# WNT8B and -9B promote the survival and dissemination of dormant ovarian cancer cells

**DOI:** 10.64898/2026.08.11.744088

**Authors:** Komila Zakirova, Daniel T. Passos, Chad Kelawan, Michael V. Roes, Rizwan Tahir, Malcolm Hill, Seung J. Kim, Matthew J. Cecchini, Marco Mura, Trevor G. Shepherd, Piru Perampalam, James I. MacDonald, Frederick A. Dick

**Affiliations:** Verspeeten Family Cancer Centre, London Health Sciences Centre Research Institute, London, Ontario, Canada, N6A 4L6; Department of Pathology and Laboratory Medicine, University of Western Ontario, London, Ontario, Canada, N6A 5C1; Interstitial Lung Disease Research Laboratory, London Health Sciences Centre Research Institute, London, Ontario, Canada, N6A 4L6; Department of Anatomy and Cell Biology, University of Western Ontario, London, Ontario, Canada, N6A 5C1; Children’s Health Research Institute, London, Ontario, Canada, N6A 4V2

**Keywords:** High-grade serous ovarian carcinoma, non-canonical Wnt signalling, cancer dormancy, cancer stem cell, CRISPR screening, metastasis

## Abstract

Cancer cell dormancy and the resultant resistance to conventional therapies present significant challenges for the successful treatment of high-grade serous ovarian cancer (HGSC). We used genome wide, and specialized sgRNA, libraries in CRISPR-based screens to identify critical cell survival mechanisms in dormancy and metastasis. Our findings demonstrate that low expression Wnt ligands WNT8B and WNT9B are essential for sustaining cell survival during prolonged dormant spheroid culture conditions. These Wnt ligands utilize non-canonical signaling to activate expression of stem cell genes such as *ALDH1A1*, *CD44* and others during spheroid dormancy. The loss of WNT8B and WNT9B reduced survival of xenografted ovarian cancer cells during early dissemination of disease that extended survival. Furthermore, treatment of WNT8B/9B deficient xenografts with carboplatin demonstrated increased sensitivity that further reduced dissemination and extended survival. These findings reveal that rare Wnt ligands can possess outsized functions in cancer pathogenesis and offer new avenues for improving treatment outcomes for HGSC through their inhibition.

## Introduction

Ovarian cancer is the eighth most common cancer in women worldwide (Lheureux et al, 2019). High-grade serous ovarian carcinoma (HGSC) accounts for 70% of these cases (Lheureux et al, 2019). It is often diagnosed late due to its asymptomatic nature and unsatisfactory early screening, resulting in a five-year survival rate of only 30% (Lheureux et al, 2019; Liberto et al, 2022). The standard treatment for advanced HGSC involves cytoreductive surgery and platinum-based chemotherapy (Atallah et al, 2023). Although most patients initially respond well, they develop platinum resistance, leading to recurrence and poor prognosis (Laurent & Liu, 2024; Lukanović et al, 2022). Despite extensive research on HGSC, new treatment approaches are only slowly emerging.

Metastasis in HGSC is largely confined to the peritoneal cavity, where multicellular clusters of cancer cells, known as spheroids, disseminate through fluid to the surrounding organs (Bowtell et al, 2015; Tadić et al, 2024). Spheroid cells paradoxically exit the cell cycle and enter dormancy, a state where residual tumor cells persist post-treatment, remaining asymptomatic for extended periods but retaining the potential to initiate disease relapse (MacDonald et al, 2017; Shepherd & Dick, 2022). Spheroids are characterized by epithelial-to-mesenchymal transition (EMT) and cancer stem cell (CSCs) properties, and these contribute to the acquisition of resistance to chemotherapy (Devenport et al, 2025; Liao et al, 2014; Velletri et al, 2021). Consequently, elucidating the survival mechanisms used by cells in dormant spheroids is crucial for developing novel therapies to combat treatment-resistant, recurrent disease.

Dissemination of HGSC from its origin in the lower abdomen is closely linked with dormancy and spheroid formation (Shepherd & Dick, 2022). This cellular state is highly dependent on EMT and cancer stem cell properties (Dongre & Weinberg, 2019; Li et al, 2023). EMT plasticity, characterized by a spectrum of states with partial epithelial to mesenchymal differentiation, as well as the reverse process of mesenchymal to epithelial transition (MET), during the different stages of HGSC progression, is thought to allow ovarian cancer cells to detach, migrate, resist cell death while suspended in ascites, and seed secondary tumors (Loret et al, 2019). Indeed, EMT-hybrid cells are shown to be more aggressive than fully differentiated cells (Klymenko et al, 2017; Strauss et al, 2011). Ascites-derived ovarian spheroids that express both E-cadherin and N-cadherin markers exhibit increased invasiveness compared to purely epithelial spheroids (Klymenko et al, 2017; Strauss et al, 2011). Additionally, EMT is thought to contribute to the acquisition of stem cell properties, marked by the expression of markers, such as *ALDH1A1* and *CD44*, which further promote tumor growth and self-renewal (Celià-Terrassa & Jolly, 2020; Frąszczak & Barczyński, 2023; Zong & Nephew, 2019). Therefore, while Wnt signaling is acknowledged for its role in EMT and cancer stem-cell renewal in various human cancers, our limited understanding of its involvement in HGSC progression highlights an important area for further research.

Wnt signaling governs various cellular processes associated with dormancy and metastasis, such as cell proliferation, polarity, survival, stem cell fate determination during embryonic development, and maintenance of homeostasis in adult tissues (Chidiac & Angers, 2023; Maurice & Angers, 2025; Steinhart & Angers, 2018). In the context of ovarian cancer, aberrant Wnt signaling has been shown to correlate with chemoresistance (Chau et al, 2013), EMT plasticity during metastasis (Arend et al, 2013; Celià-Terrassa & Jolly, 2020), and poor patient outcome (Boone et al, 2016). There are three branches of Wnt signaling including the canonical pathway that activates β-catenin, and two non-canonical pathways (Akoumianakis et al, 2022). These include the planar cell polarity pathway and the Wnt/Ca^2+^ pathway (Akoumianakis et al, 2022). There are few mutations in Wnt signaling components in the HGSC histotype of ovarian cancer (Nagaraj et al, 2015). In contrast, deregulation of the Wnt/β-catenin pathway, primarily due to mutations in the *CTNNB1* gene and, less frequently, from inactivating mutations in the *APC*, *AXIN1*, or *AXIN2* genes, is common to other gynecological malignancies such as endometrioid and mucinous ovarian cancer (Arend et al, 2013; Wu et al, 2001). Nonetheless, without recurrent genetic alterations, Wnt signaling is reported to be altered in HGSC, often as shown by nuclear or cytoplasmic localization of β-catenin (Kildal et al, 2005; Lee et al, 2003; Wang et al, 2006). Transcript analyses indicate that Wnt ligands, such as WNT2A, WNT5A, WNT7A, WNT9A, are produced by HGSC cells, tumor-associated macrophages, and cancer associated fibroblasts (Fang et al, 2024; Kotrbová et al, 2020; Piki et al, 2023; Reinartz et al, 2016). Beyond these observations, WNT7A has been shown to activate the canonical Wnt pathway in HGSC and its expression correlates with poor outcome (King et al, 2015; MacLean et al, 2016; Yoshioka et al, 2012). WNT5A and WNT11 have shown similar expression patterns as WNT7A, their expression predicts poor outcomes (Chehover et al, 2020; Xu et al, 2022), and they have been implicated in non-canonical Wnt signaling pathways (Fang et al, 2024; Jannesari-Ladani et al, 2014; Kotrbová et al, 2020; Piki et al, 2023). In contrast to this oncogenic role, Wnt signaling has also been reported to be silenced in HGSC, suggesting a tumor suppressive context as well (Kim et al, 2015; Liu et al, 2019; MacLean et al, 2016). Collectively, these studies emphasize the importance of Wnt signaling in HGSC, however, the emphasis on expression levels and correlations with outcome leaves the mechanism of Wnt participation in HGSC largely unexplored. Furthermore, it is unclear whether individual Wnt ligands are essential to HGSC pathogenesis, or if extensive redundancy indicates a collective role for the Wnt family. Of particular relevance to this report, WNT8B and WNT9B are relatively unstudied outside of roles in development (Carroll et al, 2005; Fotaki et al, 2010; Jin et al, 2020; Lan et al, 2006), and not previously linked to HGSC pathobiology.

We have performed genome-wide CRISPR screens in three HGSC cell lines to identify genes essential for cell viability in dormant spheroids (Perampalam et al, 2024a). These data indicate components of Wnt signaling are critical to the survival of spheroids under dormant culture conditions (Perampalam et al, 2024a). We selected the top-scoring essential genes from our previous study and performed a new CRISPR screen evaluating just these top scoring genes in spheroid survival. This new comparison of viability genes reveals Wnt signaling is vital for dormant spheroids, ranking them highly even in comparison with essential genes that have been previously identified. We show that the Wnt ligands, specifically WNT8B and WNT9B, whose expression in HGSC cells is low in comparison to other Wnt ligands, promote spheroid self-renewal when continually cultured under suspension conditions. Inhibiting Wnt signaling via knockout of *WNT8B* and *WNT9B* genes, or both together, resulted in reduced cell survival in dormant spheroid culture. Consistent with a self-renewal role, WNT8B/9B deficiency blocked expression of *ALDH1A1*, *CD44* and other cancer stem cell genes, as did downstream components of non-canonical Wnt signaling. Finally, our results demonstrate that WNT8B/9B signaling contributes to metastatic spread *in vivo* in xenografted mice and its loss potentiates the effect of carboplatin in culture and in xenografts. Our research highlights the critical role of WNT8B and −9B signaling in maintaining the viability of spheroids in a dormant state and in activating cancer stemness programs downstream of this pathway.

## Methods

### Cell lines and culture conditions

For gene knock out, spheroid culture, and xenograft experiments, human ovarian cancer cell lines OVCAR3, OVCAR4, OVCAR5, OVCAR8, HeyA8, TOV1946, iOvCa147 and COV318 were used. Use of iOvCa147 and TOV1946 HGSC cells for CRISPR screens has previously been reported (Perampalam et al, 2024a). HGSC cells were cultured in complete media: DMEM/F12 (Gibco CAT#11320033), supplemented with 10% FBS (Wisent CAT#090150) and 1% pen-strep-glutamine (Gibco CAT#10378016). Standard tissue culture-treated polystyrene plates (Nunc CAT#150468) were used to expand adherent cultures, while spheroid cultures were maintained on ultra-low attachment (ULA) tissue culture plates (Corning CAT#4615). All cell line identities were confirmed by STR analysis through the services of The Centre for Applied Genomics (TCAG, Toronto).

### Mini CRISPR screen in OVCAR8 cells

The mini CRISPR screen focused on 2,740 essential genes common to both OVCAR8 and iOvCa147 cell lines, identified through genome-scale CRISPR screens. The mini CRISPR library comprised a total of 17,840 sgRNAs, with 6 sgRNAs per gene from the 2740 genes; guide sequences were derived from the human GeCKOv2 lentiviral library (Sanjana et al, 2014), along with non-targeting control sgRNAs for *Luc*, *EGFP*, and *LacZ* genes. Viruses were packaged by transfecting HEK293T cells with the library and packaging components as described previously in Perampalam et al. 2024 (Perampalam et al, 2024a). 72 hours after transfection, media containing viral particles was collected, filtered through a 0.45 μm filter, and stored at −80°C with 1.1 g/100 mL BSA. OVCAR8 target cells were engineered to express Cas9 using pLentiCas9-Blast as described previously (Perampalam et al, 2024a). OVCAR8 Cas9-positive and Cas9-negative cells were transduced separately at an MOI of 0.3, such that an infected cell receives only one sgRNA. 24 hours after transduction, cells were selected with 4 μg/ml puromycin for one week. The predicted library representation was >700-fold, with over 3.75×10^7^ cells collected after selection. Cas9 positive and negative cell populations were divided into three groups, each containing 700x library coverage (approximately 1.25×10^7^ cells). Cell samples from each were frozen as pellets for genomic DNA extraction at day 0. The remaining cells were plated in 15×15 cm standard culture dishes and grown for 48 hours, after which the three replicates of 1.25×10^7^ cells each were collected and frozen for genomic DNA extraction from the adherent population. The leftover cells were seeded at 7.5×10^5^ cells/ml in 20 ml onto 10 cm ULA dishes. After 48 hours of incubation in suspension culture, media containing spheroids was transferred to 15 cm standard tissue culture plates. The following day, unattached spheroid cells were collected and re-plated onto additional 15 cm plates. This process was repeated for a total of 5 days to maximize the capture of viable cells. Attached cells were collected and pooled for DNA extraction from this re-attached spheroid population. Genomic DNA was extracted from cell pellets using the QIAamp DNA Blood Maxi Kit (Qiagen, #51194). Guide RNA inserts were amplified by PCR using NEBNext Q5 Ultra Master Mix (NEB CAT#MO544) and primers harboring i5 and i7 barcodes from Hart et al. 2015 (Hart et al, 2015). PCR amplicons were run on an ethidium-bromide stained 2% agarose gel and extracted using Monarch DNA Gel Extraction Kit (NEB CAT#T1020). DNA yields were quantified using an Illumina NextSeq 75 cycle High Output kit on an Illumina NextSeq platform using the services of the London Regional Genomics Centre (LRGC, London, Canada). Reads were mapped and count tables analyzed using TRACS (Perampalam et al, 2024b). Enrichment ratio data for all genes tested can be found in **Supplemental Table 1**.

### Spatial transcriptomics of patient derived spheroids

Spheroids were isolated from patient ascites and collected by filtration as described previously (Perampalam et al, 2024a), fixed with formalin and embedded in paraffin blocks. Acquisition of patient samples was approved by institutional REB (Project ID #115904). 6 mm cores were cut from each patient block and embedded in a new paraffin block to cluster spheroids from three patients into an 11mm capture area of a 10X Genomics Visium slide. Tissue was captured according to the manufacturer’s protocol (CG000495, Rev E, 10X Genomics). High resolution tissue scans were taken with a Grundium Ocus 20. Samples were sequenced at the Princess Margaret Cancer Centre Genomics facility (Toronto). Using 10X Genomics Space Ranger Count command (3.1.2 or 4.0.1), reads were aligned to the GRCh38 human transcriptome and Visium human transcriptome Probe Set v2.1.0 (GRCh38-2024-A) references and mapped to tissue spots with microscope images of the Visium slides for barcode and UMI mapping. Tissue and fiducial frame detection of tissue images were manually performed using the 10X Genomics Loupe Browser Visium Manual Alignment tool. Gene expression of barcoded spots within tissues was subsequently analyzed using 10X Genomics Loupe Browser software. Tissue spots were filtered by setting a minimum threshold of 100 UMIs and 100 genes per barcode. Tissue spots relating to specific cell lines or patient samples were manually annotated to compare gene expression patterns of whole tissue samples.

### Generation of gene knockout cell lines

Multiple sgRNA sequences were used to target *WNT8B*, *WNT9B*, *APC*, and *PCDH8* genes and they were derived from the GeCKOv2 library (Sanjana et al, 2014). See Supplemental Table 2 for sgRNA sequences used to make these vectors. The negative control cells were created using sgRNAs targeting *LacZ*, *Luc*, and/or *EGFP* genes. Oligonucleotides containing sgRNA sequences were synthesized by Thermo Fisher and cloned using Gibson Assembly (NEB CAT#E2611) as described (Wang et al, 2016), or BbsI and Golden Gate Assembly as reported (Kabadi et al, 2014). All plasmid constructs were confirmed by Sanger sequencing relevant regions (LRGC, London, Canada). The resulting plasmid or plasmid pool was transfected into HEK293T cells along with plasmids encoding lentiviral packaging proteins (Perampalam et al, 2024a). After 48 to 72 hours, media containing viral particles was collected, centrifuged at 500 rcf to pellet dead cell debris, and then filtered using a 0.45 μM PES syringe filter (Sarstedt CAT# NC9410831). All cells were transduced at a high MOI to increase the chances that each target cell received a full complement of guides. One day following transduction, cells were selected with 4 μg/ml puromycin for 72 hours or used immediately.

### Spheroid culture, reattachment viability assays, and western blotting

A total of 3-6 × 10^5^ HGSC cells were seeded in 6-well flat-bottom ULA plates (Corning CAT#3471) to facilitate spheroid formation. Controls and knock outs were always compared at equal seeding density. Spheroids were trypsinized every 7 days, resuspended in complete media, and replated onto ULA plates to produce successive generations. Microscopy images of spheroids were captured at the end of each week-long generation, over three consecutive generations. At the end of the 21-day experiment, spheroids were reattached for 24 to 48 hours in standard 6-well tissue culture plates (Nunc CAT#140675) to isolate viable cells. To assess sensitivity to Carboplatin, an equal number of cells (6 × 10^5^) of OVCAR8 sgRNA control or WNT DKO were treated with a single dose of different Carboplatin concentrations (1 μM, 2.5 μM, 5 μM, or 10 μM) and incubated for 7 days in suspension culture conditions without media change. Saline was used as the vehicle control.

Crystal violet staining was performed to compare cell abundance among different cell genotypes or drug treatment conditions. Cells were first fixed in 100% methanol for 10 minutes, then shaken for 30 minutes in a solution of 0.5% crystal violet and 25% methanol in PBS. Plates were gently submerged in water to remove any residual crystal violet and then allowed to dry. Next, the wells were destained by adding 1 ml of 10% acetic acid in PBS and shaking for 1 hour to extract the crystal violet from the stained cells. Crystal violet absorbance was measured at 590 nm using a microplate reader (Perkin Elmer Wallac Victor2 1420). Comparison of this data with Trypan blue dye exclusion and CellTiter-Glo methods confirms that the absorbance reading of Crystal Violet in this assay is directly proportional to the number of reattached cells and is presented as relative viability in this study (MacDonald et al, 2017; Perampalam et al, 2024a). For western blotting experiments cells were washed with cold PBS and lysed in RIPA buffer. Equal amounts of protein were loaded in SDS-PAGE gel lanes and blotted using standard methods. Sources of antibodies and dilutions used can be found in the key resources listed in **Supplemental Table 3**.

### Verification of the double knockout genotype for WNT8B and WNT9B in OVCAR8 cells and real time RT-PCR

*WNT* DKO cells were generated by transducing OVCAR8 *WNT8B* individual knockout cells with lentiviruses targeting the *WNT9B* gene. Puromycin selection was not performed after the second round of lentiviral infection, as *WNT8B* knockout target cells had already acquired resistance from the prior infection. The double knockout genotype of the heterogeneous population of OVCAR8 cells was confirmed by extracting RNA using the Monarch Total RNA Miniprep Kit (NEB CAT#T2010S), converting RNA to cDNA, and conducting RT-qPCR analysis of *WNT8B* and *WN9B* mRNA levels using syber green master mix (BioRad) (See **Supplemental Table 2** for all PCR primers).

Additionally, 5 monoclonal *WNT* DKO cell lines were isolated through limiting dilutions of the heterogeneous pool of OVCAR8 *WNT* DKO cells, which were then expanded for genomic DNA extraction, PCR amplification of putative sgRNA target sites, and Sanger sequencing. The bioinformatics-based tool COSMID (CRISPR Off-target Sites with Mismatches, Insertions, and Deletions) was used to search for potential off-target sites in the human genome as described in Cradick et al., 2014 (Cradick et al, 2014). On-target exon regions of both *WNT8B* and *WNT9B* genes, along with the highest-ranked potential off-target genomic locations were evaluated for Cas9-mediated mismatches. Using primers generated by COSMID, genomic locations of interest were amplified by PCR. The DNA fragments were then gel-purified using the Monarch DNA Gel Extraction Kit (NEB CAT#T1020) and cloned into pJET1.2/blunt vector from the CloneJET PCR Cloning Kit (Thermo Scientific, CAT# K1231). The ligation mixture was directly used for transformation into *E. coli* DH10B competent cells (Thermo Scientific CAT# FEREC0113). 5 to 10 bacterial colonies per each PCR ligation mix were selected and inoculated into LB broth containing 100 µg/mL ampicillin for overnight culture at 37 °C with shaking. Plasmids were purified using Monarch Plasmid Miniprep Kit (NEB CAT#T1010) and sent for Sanger sequencing (LRGC, London, Canada) using the pJET1.2 forward and reverse sequencing primers. DNA sequences were aligned with the reference GRCh38.p14 human genome using SnapGene to determine the consequences of indels that were detected.

### Time-lapse experiments using fluorescent reporter constructs

The canonical Wnt lentiviral reporter construct pBarVenus (Biechele & Moon, 2008) was provided by the Angers lab, which contains a concatemer of 12 synthetic TCF-binding sites driving the expression of Venus. The stemness reporter vector was engineered by excising *TCF*-binding sequences with PaqCI and XmaI restriction enzymes and substituting them with a 1kb proximal promoter region from *ALDH1A1* using Gibson Assembly (NEB CAT#E2611). The *ALDH1A1*-Venus reporter vector was additionally modified to confer resistance to neomycin through the replacement of puromycin cassette, facilitating the selection of OVCAR8 WNT DKO cells that are already resistant to puromycin. OVCAR8 negative control and *WNT* DKO derivatives were separately transduced with reporter constructs, followed by treatment with 400 μg/ml G418 for a duration of 10 to 14 days.

A total of 2,000 cells were seeded into each well of round-bottom 96-well ULA plates (Corning, Cat# 7007) in 200 μL of complete media. To activate canonical Wnt signaling, 1 μM CHIR99021 (Stemcell, CAT#72054) GSK3 inhibitor was added per well immediately after seeding cells into suspension culture. The plate was placed in the drawer of IncuCyte S3 Live-Cell Analysis System (Sartorius) for the time-lapse analysis. Brightfield and fluorescent images were captured at 3-hour intervals over a period of 7 days. The acquisition time for the green channel was set to a default of 300 ms. Changes in total integrated intensities over a week were plotted using GraphPad Prism.

### Immunofluorescence staining of OVCAR8 cells and derivatives

Glass slides were pretreated with poly-D-lysine (Gibco CAT#A3890401) and left to dry in the incubator overnight at 37°C and 5% CO_2_. Glass slides were placed in the wells of a 6-well plate, with a total of 2×10^5^ cells plated on each slide in complete media and allowed to attach. 24 to 36 hours after cell seeding, the media was aspirated, and cells were washed three times with 1x PBS for 5 minutes each on a shaker. Cells were fixed with 4% paraformaldehyde (pH 7.4) for 10 minutes at room temperature. 400 µL of 0.3% Triton X-100 in 1x PBS was added for 10 minutes to permeabilize the cells. 0.3% Triton X-100 + 5% BSA in 1x PBS was used as the blocking buffer for 1-2 hours at room temperature. Primary antibodies were diluted in 1xPBS + 0.3% Triton X-100 + 5% goat serum as follows: rabbit polyclonal anti-β-catenin antibody (Abcam CAT#ab6302) 1:2000 dilution; mouse monoclonal anti-APC antibody (Abcam CAT#ab16794) 1:1000 dilution. The secondary antibodies used: Alexa Fluor 594 goat-anti-mouse IgG (Invitrogen CAT#A-11005); Alexa Fluor 594 goat-anti-rabbit IgG (Invitrogen CAT#A-11012) diluted 1:800 in blocking buffer and incubated for 1-2 hours. A drop of ProLong Gold Antifade Mountant with DNA Stain DAPI (Thermo Fisher CAT#P36935) was used for nuclear counterstaining. For negative controls, glass slides were stained only with secondary antibodies or with DAPI counterstain. Slides were mounted on double-frosted microscope slides (Fisherbrand CAT#12-552-5) with SlowFade Diamond Antifade Mountant (Thermo Fisher CAT# S36967). Fluorescence images were captured using Leica DMI6000B inverted microscope.

### Xenografts in immunocompromised mice

We used ARRIVE principles to guide the design of our xenograft experiments. All procedures adhered to our animal protocol approved by the Western Animal Care Committee (protocol 2024-023). Female NOD.Cg-Prkdc^scid^/J mice (JAX CAT#001303), were randomly selected between 6 to 8 weeks of age for intraperitoneal injections of 4×10^6^ OVCAR8 negative control or WNT DKO derivatives. To model the treatment of residual disease, xenografted mice were again randomly selected for intraperitoneal Carboplatin injections at a dose of 10 mg/kg or with 0.09% saline vehicle control. First treatments were administered on the same day as cell engraftment, and one week later, for a total of two doses, irrespective of the length of the experiment. At 14- and 35-days post-engraftment, mice were euthanized and necropsied to assess disease spread. Abdominal cavities of mice were rinsed with sterile PBS to collect disseminated spheroids, and a blinded lab member seeded them into 6-well ULA plates containing complete media. They were subsequently incubated for 72 to 120 hours in suspension culture. For viability comparisons, spheroids were further reattached to 6-well standard culture plates and stained with Crystal Violet as outlined above.

### Alu-based qPCR quantification of human cancer cells

Genomic DNA was extracted from cells collected via abdominal PBS washes using the Monarch Genomic DNA Purification Kit (NEB #T3010). For precise comparisons of cancer cell dissemination among mouse groups, genomic DNA was extracted from equal volumes of PBS washes and then eluted in a consistent final volume of 100 µl of nuclease-free water per tube. To enhance the yield of genomic DNA, carrier RNA (Sigma-Aldrich CAT#GE27-4110-02) was added to the Binding Buffer at a concentration of 10 µg/ml. 1 µL of the isolated genomic DNA was used for qPCR of human Alu repeats, using primers (101F and 206R) and probe (144RH) from Funakoshi et al. 2017(Funakoshi et al, 2017). PCR was performed in a total volume of 20µl with TaqMan Universal Master Mix II, no UNG (Thermo Fisher Scientific). Technical duplicates were processed in a single run, and the mean Ct values were used to determine the quantity of human cells in every sample. To quantify human cell numbers from genomic DNA amounts, we generated a standard curve using known concentrations of human genomic DNA that were serially diluted in mouse genomic DNA to establish standards and detection limits.

### Data and statistical analysis

CRISPR screens were analyzed using TRACS (Perampalam et al. 2024)(Perampalam et al, 2024b).

TCGA HGSC cancer genomic data was queried using cBioPortal (Bell et al, 2011; Heath et al, 2021).

Statistical analysis was performed in GraphPad Prism (Version 10).

Diagrams and schematics were generated in Biorender.com

DNA sequences alignments were analyzed in SnapGene (Version 8).

### Data availability

The CRISPR screen sequencing data generated in this study are publicly available in Gene Expression Omnibus (GEO) at GSE306119. Spatial transcriptomic sequencing data is available at GSE335181.

## Results

### The Wnt pathway contributes to the survival of dormant HGSC cells

The formation of non-proliferative HGSC spheroids in suspension culture is a stressful process during which many cells die without being incorporated into a spheroid. To identify the survival genes whose loss compromises viability during dormancy, we developed a genome-wide CRISPR screening method that addresses this biological challenge. It integrates data from both actively proliferating cells grown in standard monolayer (2D) and cells cultured in three-dimensional (3D) spheroid conditions, mimicking the formation of dormant aggregates found in the ascites of ovarian cancer patients. The methodology for our genome-wide CRISPR screening approach has been described previously (Perampalam et al, 2024a; Perampalam et al, 2024b). **Figure 1A** illustrates a simplified overview of the screens conducted across three HGSC cell lines, OVCAR8, iOvCa147, and TOV1946, which uncovered 1,382 genes that are commonly essential for survival in dormancy in these cell lines (Perampalam et al, 2024a; Perampalam et al, 2024b). Pathway analysis of these 1,382 common genes identified Wnt signaling as one of the most enriched secondary Reactome categories (**Figure 1B**). Enrichment Ratio (ER) values denote the log_2_-fold change in the ratio of gene scores between spheroid and adherent conditions. The genes with a log_2_ ER value less than 0 (p_adj_<0.05) are scored as essential for dormancy, as their loss impairs cell survival in suspension culture more than in adherent, proliferative culture conditions. **Figure 1C** shows ER and P-values for all individual WNT genes in the screened cell lines as bubble plots indicating that a few are required for survival in dormant culture conditions in multiple cell lines. A mean ER for each WNT gene deletion was also calculated to illustrate the collective functional consequence of deletion of these ligands in dormancy (**Figure 1D**). This data is accompanied by mean gene effect scores for all WNT ligands from DepMap HGSC data that shows the very modest effects of deleting these genes in adherent, proliferating cells. Lastly, we examined the expression levels of all WNT ligands in HGSC from TCGA data (**Figure 1E**, left), and quantitated the induction of expression of WNTs identified in our screen or previously reported to impact HGSC pathobiology (**Figure 1E**, right). This data demonstrates a genetic dependency for WNT ligands in dormant cell culture conditions that is distinct from proliferating cells (eg. WNT8B, −9B, and −10A). Furthermore, genetic dependency in dormancy is observed for some of the weakest expressed WNTs (WNT8B and −9B), including family members that are uninduced in dormancy (WNT9B). This view of WNT family members highlights novel ligands WNT8B, −9B, and −10A as having the greatest dependency in HGSC dormant culture conditions.

**Figure 1.**
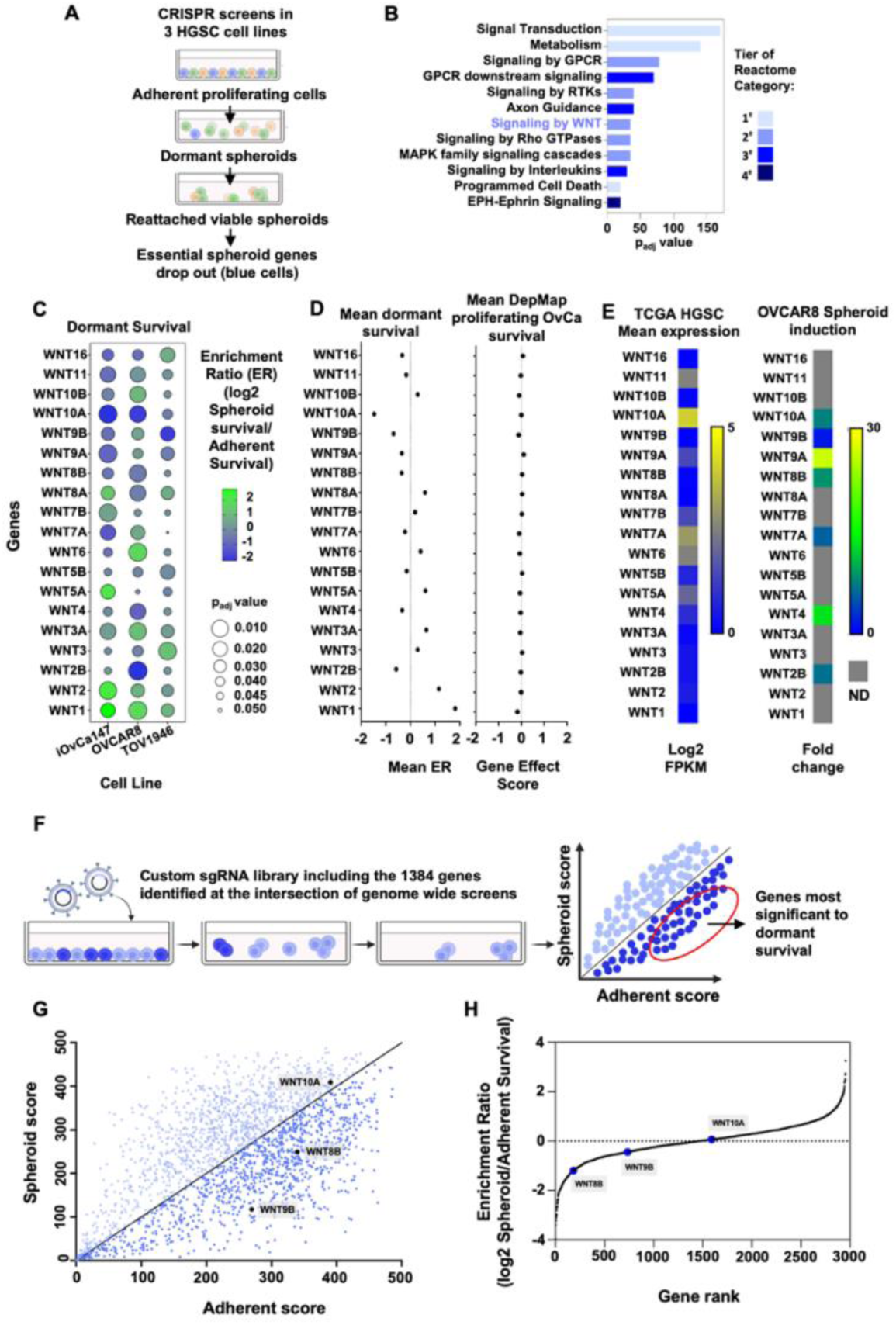
CRISPR screens identify the Wnt signaling pathway in HGSC dormant cell survival. **(A)** Flow chart of genome-wide CRISPR screens conducted in each of iOvCa147, TOV1946, and OVCAR8 cell lines. The screen identified genes whose function supports viability in the dormant spheroid step of this culture procedure. **(B)** Gene ontology analysis conducted on the 1382 shared genes identified in A. Tiers of Reactome categories are indicated by shading. **(C)** Bubble plots illustrate the effect on cell survival of mutations in each WNT ligand gene in each cell line in suspension culture. Genes with a log_2_ Enrichment Ratio (ER) <0 are shaded in blue, with darker blue indicating a smaller ER and a greater impact on cell survival during dormancy. The size of the bubbles reflects p_adj_ value for the indicated gene. **(D)** Graph showing the mean ER for each of the indicated WNT ligand genes across all three cell lines. For comparison, the mean gene effect score for each WNT ligand derived from DepMap HGSC cells under adherent proliferating conditions is shown. **(E)** Heat map depicting relative gene expression of each WNT ligand from TCGA HGSC RNA-seq data. For comparison, qPCR was used to quantitate the fold change in expression of the indicated WNT ligands when adherent OVCAR8 are transferred to suspension culture for seven days. ND=not determined. **(F)** Flow chart of mini CRISPR screen conducted in OVCAR8 cell line. The genes with log_2_ ER <0, which were commonly identified in genome-wide CRISPR screens across the three HGSOC cell lines, were targeted by a custom sgRNA lentiviral library. In the mini CRISPR screen, these essential genes were compared to each other, with light and dark blue shading indicating lower or higher effects on cell viability in dormant suspension cultures. The spheroid score (y-axis) and the adherent score (x-axis) for each gene, which were then plotted as a scatter plot. **(G)** The mini CRISPR screen results are shown with relevant WNT ligands labeled. **(H)** Gene rank plot to illustrate WNT ligand relative efffects in this screen.

We constructed an sgRNA library by selecting genes with a log_2_ ER below 1 that were commonly identified in our three genome-wide screens and performed an additional CRISPR screen to compare these essential genes amongst themselves. This library contained sgRNAs targeting 2,740 genes and non-targeting control guides for *LacZ*, *Luc*, and *EGFP*. This list incorporates the 1,382 genes shared among the three HGSC cell lines screened, as well as those found at the intersection of OVCAR8 and iOvCa147, including *WNT8B*, *WNT9B*, and *WNT10A*. This CRISPR screen was conducted in OVCAR8 cells following the previously outlined experimental workflow (Perampalam et al, 2024a; Perampalam et al, 2024b) (**Figure 1F**), where dark blue indicates the greatest negative impact on cell viability from a gene loss event in dormant culture. Our findings highlight WNT ligands, particularly *WNT8B* and *WNT9B*, as genes with the low ER values, indicating that their loss is more harmful to HGSC cells in spheroid culture, even among genes already deemed essential during dormancy (**Figure 1G, H**). This discovery further implicates novel HGSC WNTs, −8B and −9B in dormant spheroid cell survival.

### WNT8B and 9B are essential for spheroid formation and cell survival

We next examined the functional significance of Wnt signaling through WNT8B and WNT9B ligands in spheroid survival as they have not previously been implicated in HGSC. We utilized spheroids isolated directly from HGSC patient ascites and embedded in paraffin blocks to investigate gene expression using the 10X Genomics Visium spatial transcriptomic platform (**Figure 2A**). This analysis confirmed that low level, but detectable expression of both WNT8B and WNT9B are present in dormant spheroids in these three patients (**Figure 2B**). We generated single and double gene knockouts of *WNT8B* and *WNT9B* in OVCAR8 cells (**Supplemental Figure 1A**). Successful gene disruption was confirmed by RT-qPCR analysis that demonstrated only minimal expression of the mRNAs encoding these genes in targeted cells (**Supplemental Figure 1B**). Similarly, Sanger sequencing of the targeted genomic DNA regions from several single-cell clones, confirmed that no functional alleles of *WNT8B* or *WNT9B* were found in the knockout cells and the most likely off target locations remained unaltered (**Supplemental Figure 1C**). OVCAR8 cells deficient for either of these Wnt ligands, or both (referred to as *WNT* DKO), were cultured in suspension to establish successive spheroid generations. In this experiment, enzymatic dissociation was used every 7 days followed by suspension culture and re-formation of spheroids. Spheroids were passaged for three generations with a total incubation period of 21 days in suspension culture, after which they were transferred to adherent culture plates to isolate viable cells by re-attachment (**Figure 2C**). Microscopic images of spheroids in culture show that, compared to individual *WNT8B* or *WNT9B* knockout cells, OVCAR8 non-targeting sgRNA control cells formed larger and more compact spheroids during the first seven days (**Figure 2D**). For all genotypes of cells, floating cells and debris were observed to increase with prolonged passaging in suspension culture, suggesting cell death. Notably, this was more pronounced in individual *WNT8B* or *WNT9B* knockout cells, and it was even more severe in double knockout cells (**Figure 2D**). We observed fewer and smaller spheroids at the end of 21 days in suspension culture, and the reduction in viable cells was confirmed by reattachment and staining (**Figure 2E, F**). Our results indicate that WNT8B and WNT9B function to preserve viability over time in suspension culture and confirm their discovery in our CRISPR screens. In addition, we performed time-lapse video microscopy analysis of early spheroid formation, immediately after seeding cells into suspension conditions (**Supplemental Figure 2A**). It showed that OVCAR8 control cells assemble into a spheroid within 6 hours in suspension culture. In contrast, the *WNT8B* and *WNT9B* double knockout cells do not form a compact spheroid even 24 hours post-seeding (**Supplemental Figure 2B**). These findings suggest that WNT8B and WNT9B signaling is crucial not only for the long-term self-maintenance of HGSC cells during dormancy, but also for the clustering of these cells into a dense spheroid. We similarly generated WNT8B and −9B DKO cells in six additional HGSC cell lines. These cells were placed in suspension culture to induce dormancy and spheroid formation. Following seven days of culture photomicrographs were taken to characterize spheroid formation (**Figure 2G and Supplemental Figure 2C**). This analysis demonstrated a dramatic loss of spheroid formation and cell survival in Hey A8 and iOvCa147 cell lines in just seven days. While OCVAR3, −4, −5, and COV318 formed spheroids during this time (**Supplemental Figure 2C**), only OVCAR4 maintained full viability over seven days (**Figure 2H**). This confirms a critical role for WNT8B and −9B in dormant culture across a panel of ovarian cancer cells arguing their broad importance. OVCAR8 cells show the most resilience in the absence of these two WNTs and were utilized for all subsequent experiments.

**Figure 2.**
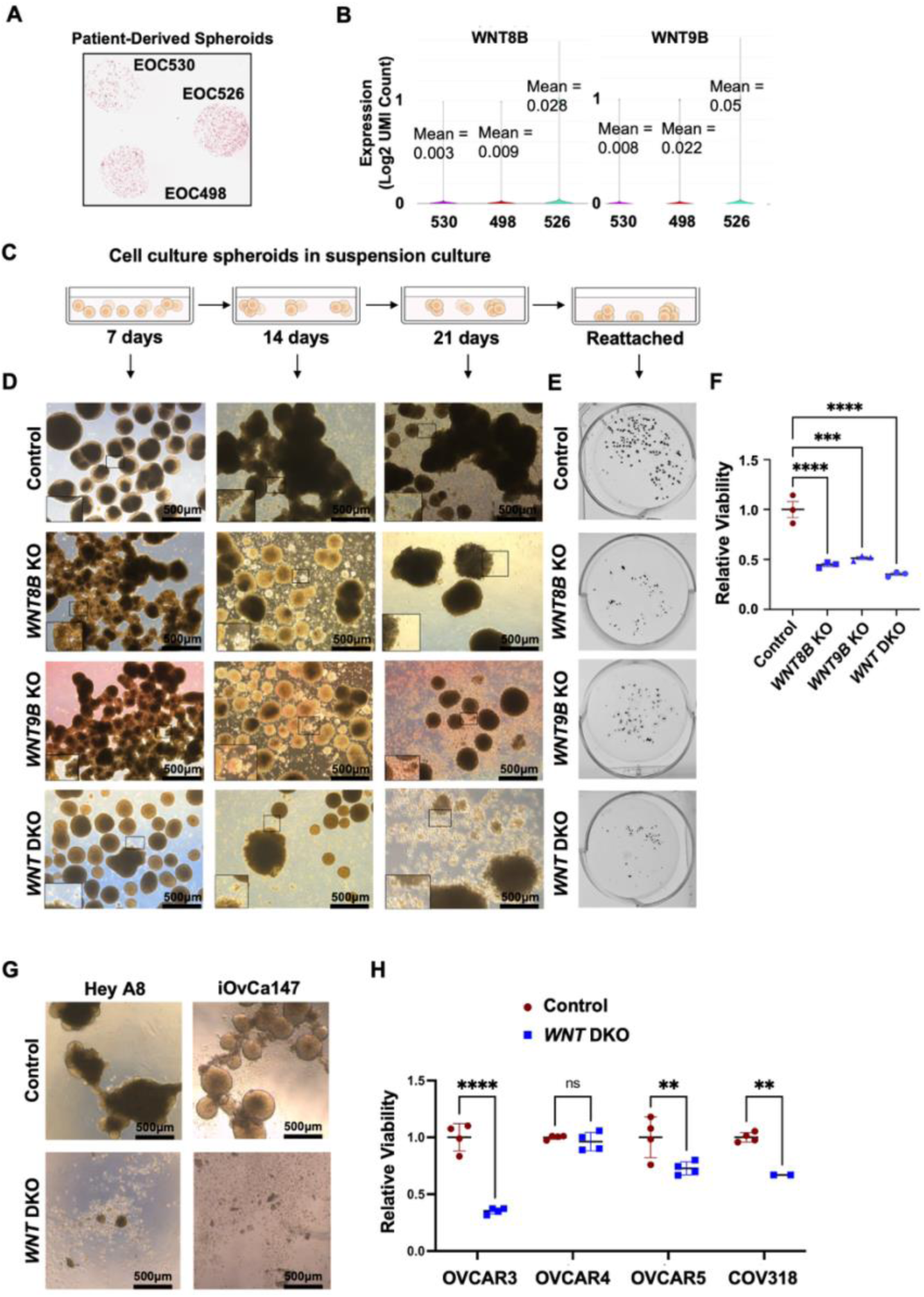
WNT8B and WNT9B ligands support spheroid formation and HGSC cell survival during dormant cell culture. **(A)** H&E staining of spheroids from three HGSC patients prior to capture and sequencing with 10X Genomics Visium. **(B)** Average gene expression of WNT8B and WNT9B is shown as violin plots with mean log2 UMI values listed for each patient. **(C)** A schematic diagram illustrating the establishment of successive spheroid generations. Spheroids were trypsinized every 7 days and incubated in suspension culture for a total of 21 days, followed by the reattachment of viable cells in adherent culture plates. **(D)** Microscopic images of spheroids were taken at 4X magnification every 7 days before passaging. Magnifying boxes zoom in on peripheral zones of spheroids with increased cell debris. **(E)** After 21 days in suspension culture, spheroids were reattached to adherent culture plates and stained with Crystal Violet dye. **(F)** The absorbance of the extracted dye was measured spectrophotometrically to quantify biomass, and the averages were compared using one-way ANOVA (n=3, *** p<0.001, **** p<0.0001, error bars are ±1 SEM). **(G)** WNT8B and WNT9B DKO derivatives were created for each of the indicated HGSC cell lines. Cells were cultured under spheroid conditions for 7 days and photographed at 4X magnification. **(H)** Relative viability of spheroids from the indicated cell lines following seven days of suspension culture. Average values were compared using one-way ANOVA (n=3, *** p<0.001, **** p<0.0001, error bars are ±1 SEM).

### Non-canonical WNT8B and WNT9B signaling in dormant HGSC cells promotes cancer stemness

Both canonical and non-canonical Wnt signaling pathways have been proposed to enhance HGSC cancer pathogenesis, enabling cells to gain stem cell characteristics and EMT plasticity, which ultimately drive more aggressive disease (Alizadeh et al, 2025). To gain insights into the mechanism by which WNT8B and WNT9B promote survival under dormant conditions, we examined EMT and cancer stem cell genes in dormant suspension culture in single knock out and WNT DKO cells.

We investigated regulatory targets of WNT signaling in EMT for their ER scores in our genome wide OVCAR8 CRISPR screen (**Supplemental Figure 3A**). Most regulatory targets involved in EMT had negative log_2_ ER scores with varying levels of significance (**Supplemental Figure 3A**). Consequently, we examined the expression of mesenchymal markers Vimentin and N-cadherin, as well as the epithelial markers Cytokeratin-7 and E-cadherin in adherent proliferating (**Supplemental Figure 3B**) and dormant spheroid cells (**Supplemental Figure 3C**). Consistent with previous reports for OVCAR8 cells (Chirshev et al, 2020; Hojo et al, 2018; Russo et al, 2024), we found that adherent and spheroid cells co-expressed mesenchymal and epithelial markers. Loss of *WNT8B* and *WNT9B* resulted in reduced expression in spheroids of the epithelial marker E-cadherin, but also of the mesenchymal marker N-cadherin, with no change in either vimentin or cytokeratin expression. This suggests that the hybrid state of these cells is not shifted toward either epithelial or mesenchymal states by the loss of *WNT8B* and *WNT9B*.

Known stem cell transcriptional targets of Wnt signaling mostly have positive log_2_ ER scores, however, specific targets appeared essential for survival in dormant suspension culture of OVCAR8 cells including *ALDH1A1*, *-1A3*, *CD44*, *NOTCH1*, and *SOX9* (**Figure 3A**). Although not all reached statistical confidence levels below P=0.05. Spheroid cells that were knocked out for either *WNT8B*, *WNT9B*, or both exhibited a reduction of over 50% in the expression of *ALDH1A1, CD44, NOTCH1,* and *SOX9* stem cell associated genes (**Figure 3B**). *ALDH1A1, ALDH1A3,* and *SOX9* expression displayed further decrease in *WNT* DKO cells relative to single knockouts. Collectively, these findings indicate that WNT8B and WNT9B signaling contribute to expression of a cancer stem cell gene program essential for survival. Examination of expression of this panel of genes in spatial transcriptomic data from HGSC patient derived spheroids indicates that these genes are robustly expressed (**Figure 3C**). These findings suggest that in dormant conditions stem cell genes are expressed to support the survival of cancer cells in a WNT8B and −9B dependent manner.

**Figure 3.**
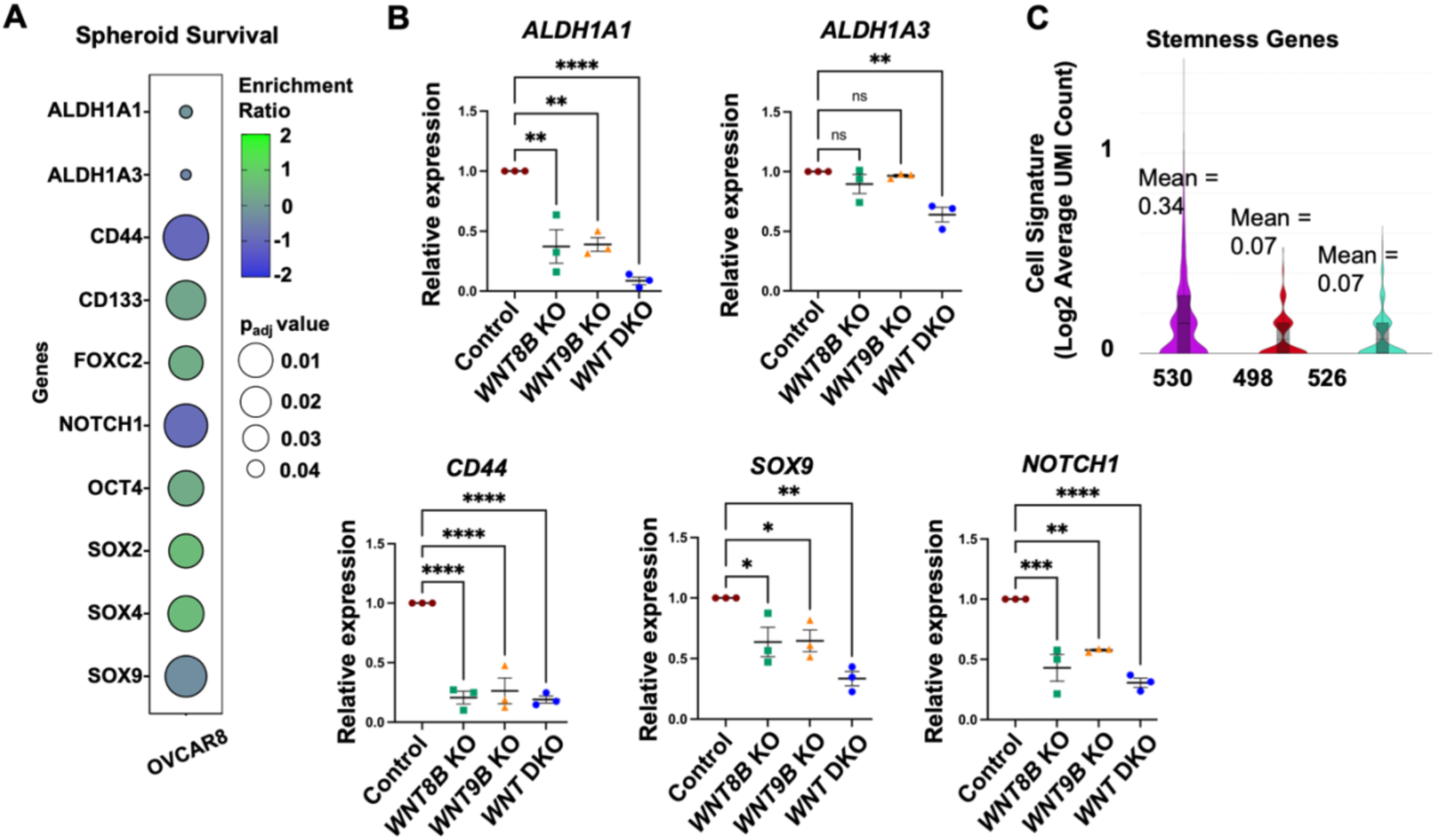
Mutation of WNT8B and WNT9B compromise stem cell gene expression in spheroids. **(A)** Bubble plots showing log_2_ ER and p-values for stem cell associated genes from the genome wide CRISPR screen in OVCAR8 cells. **(B)** Relative expression of the indicated stem cell genes was determined in spheroids from control, WNT8B, WNT9B, or DKO genotypes. Data from three to six biological samples are shown. Means were compared using one-way ANOVA (*p<0.05, **p<0.01, ***p<0.001, ****p<0.0001, error bars are ±1 SEM). **(C)** The five stem cell genes from B were used to create a gene signature and their expression in spatial transcriptomic data from patients was investigated. Average gene expression of the signature is shown as violin plots with mean log2 UMI values listed for each patient.

To examine WNT8B and WNT9B signaling specifically during spheroid formation and survival, we created a reporter that fuses 1 kb of proximal promoter from *ALDH1A1* to a Venus fluorescent reporter. We utilized lentiviral delivery to stably introduce this reporter into *WNT* DKO or a control sgRNA derivative of OVCAR8. Time-lapse imaging was performed to analyze spheroids over a 7-day period and it showed increased fluorescence from the reporter over the first three days that is maintained over the remaining four days (**Figure 4A, B**). Importantly, quantification of the fluorescence signal intensity revealed that the reporter is activated as a function of spheroid formation in dormancy and that in *WNT* DKO cells it was significantly reduced compared to OVCAR8 non-targeting control cells (**Figure 4C**). These results suggest WNT8B and −9B control cancer stem cell properties through transcriptional activation of known stem cell genes.

**Figure 4.**
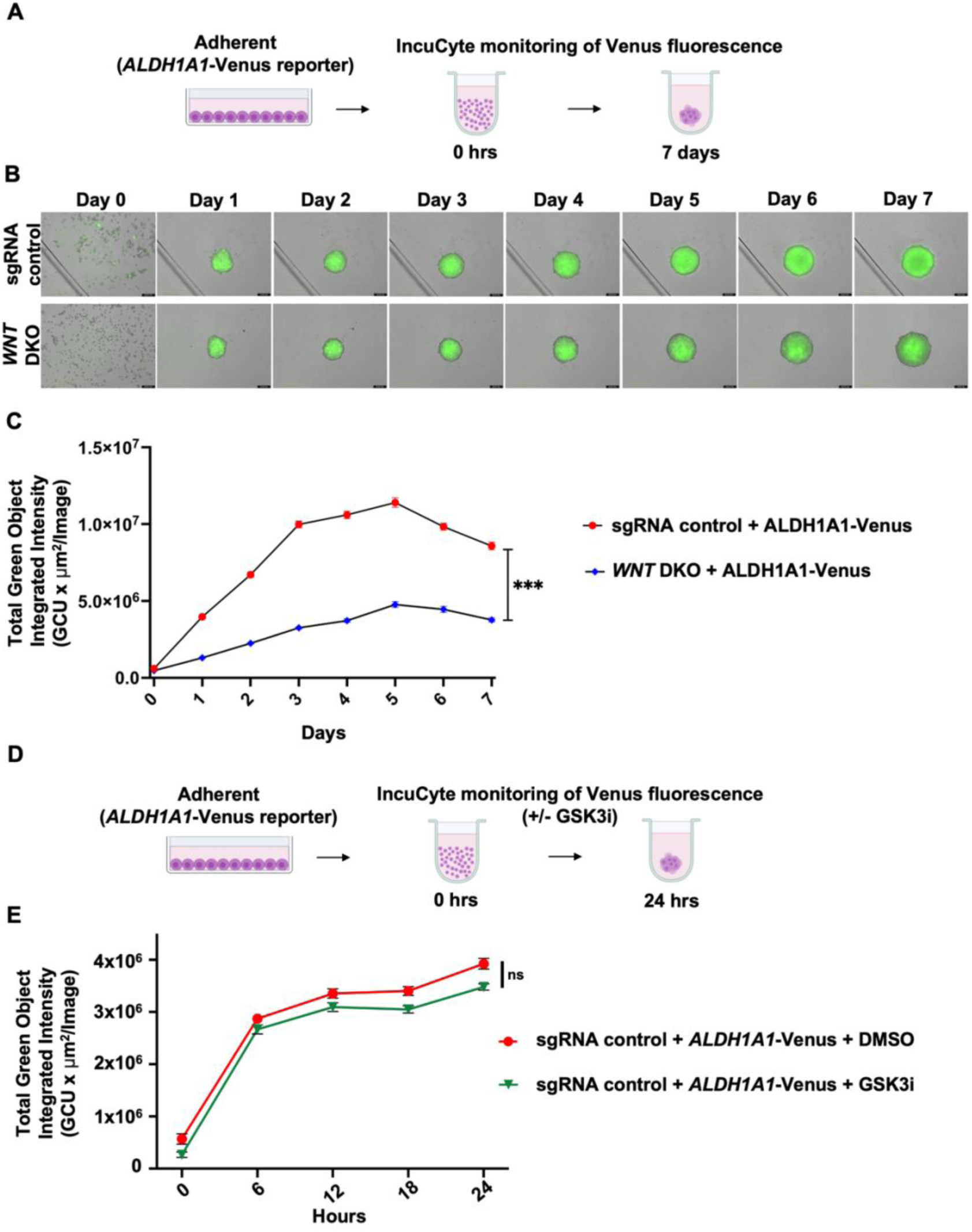
WNT8B and WNT9B activate ALDH1A1 expression in spheroids. **(A)** OVCAR8 sgRNA non-targeting control and *WNT* DKO cells were transduced with a *ALDH1A1*-Venus reporter, cultured in suspension, and fluorescent images were captured over 7 days using video microscopy. **(B)** Representative images of Venus fluorescence from spheroids are shown from immediately after seeding up to 7 days. **(C)** Integrated fluorescence intensity of Venus was quantified using the analysis software from the IncuCyte® S3 System. Statistical comparisons were made using t-test (n=6, ***p<0.001, error bars are ±1 SEM). **(D)** Cells bearing the *ALDH1A1*-Venus reporter were treated with 1 μM CHIR99021 to block GSK3 and activate canonical WNT signaling while being monitored by video microscopy. **(E)** The intensity of Venus fluorescence was measured in OVCAR8 sgRNA non-targeting control cells following treatment with either GSK3 inhibitor or vehicle DMSO for 24 hours (n=6, p=ns determined by t-test, error bars are ±1 SEM).

WNT8B and WNT9B have been reported to activate canonical signaling in development (Carroll et al, 2005; Fotaki et al, 2010; Jin et al, 2020; Lan et al, 2006), therefore we investigated the state of canonical signaling in dormant OVCAR8 spheroids. To investigate whether *ALDH1A1*-driven fluorescence is enhanced by increased canonical Wnt signaling, we treated OVCAR8 sgRNA control cells with a GSK3 inhibitor CHIR99021, which stabilizes β-catenin and activates canonical WNT signaling (**Figure 4D**). No significant difference in fluorescence intensity was observed between OVCAR8 sgRNA control cells treated with the GSK3 inhibitor and the same cells administered with vehicle only (**Figure 4E**). We further tested the status of canonical signaling in dormant spheroids through β-catenin localization in adherent culture and in spheroids and we created *APC* knock out cells to activate canonical signaling (**Supplemental Figure 4**). β-catenin was constitutively nuclear, even in spheroids (**Supplemental Figure 4A**), while APC deficiency had no effect on this status, or on viability (**Supplemental Figure 4C-F**).

To understand the nature of WNT8B and −9B signaling in spheroids, we investigated the status of both canonical and non-canonical WNT pathway components from our genome wide CRISPR screen. We note that canonical signaling components are largely characterized by positive log_2_ ER scores, suggesting they are dispensable in spheroids (**Figure 5A**). In contrast, some components of non-canonical pathways appear as strong screen hits (**Figure 5A**). We compared expression and activation of WNT receptors in adherent, proliferating OVCAR8 cells with dormant, suspension culture. **Figure 5B** shows LRP6 activation status by western blotting and detection of phosphorylation dependent migration shifts. In contrast to WNT3A activation of canonical signaling in HEK293 cells that increases the LRP6 shift, OVCAR8 cells showed little evidence of this shift in adherent culture conditions and LRP6 receptor expression appeared to diminish in suspension culture. Conversely, the non-canonical receptor ROR2 appears to increase in expression in spheroid conditions. This suggests that receptor composition is altered to favor non-canonical signaling in spheroids. Using our screen data from **Figure 5A** as a guide, we selected non-redundant genes that encode components of the planar cell polarity and calcium pathways of non-canonical Wnt signaling to test their role in regulating stem cell gene expression in spheroids (**Figure 5C**). Knocking out components of either arm of non-canonical Wnt signaling blocked *ALDH1A1*, *ALDH1A3*, and *SOX9* expression in spheroids (**Figure 5D**). This suggests that WNT8B and −9B control stem cell gene transcription through non-canonical signaling in HGSC dormant spheroids.

**Figure 5.**
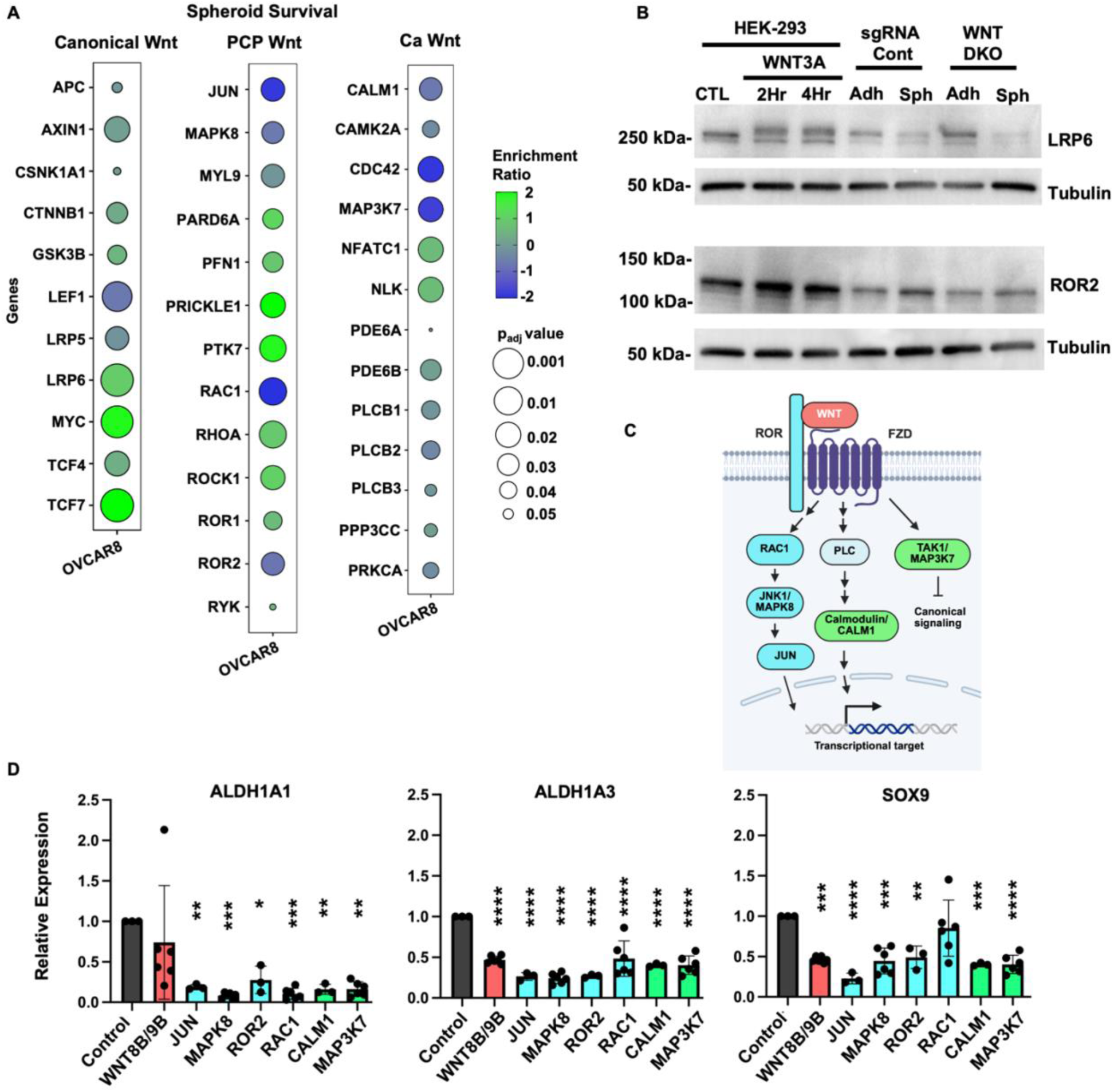
WNT8B and WNT9B utilize non-canonical signaling to activate stem cell gene expression. **(A)** Bubble plots showing log_2_ ER and p-values for select canonical WNT signaling pathway genes and non-canonical PCP and Ca pathways components from the genome wide CRISPR screen in OVCAR8 cells. **(B)** Protein extracts from the indicated genotypes of cells and culture conditions were blotted for WNT receptors LRP6 and ROR2. The relevant migration positions were labeled, and α-Tubulin was blotted as a loading control. WNT3A conditioned media was used to stimulate HEK-293 cells to illustrate effects of canonical signaling on WNT receptors. **(C)** Schematic of non-canonical pathways and the components that were deleted to test their role in regulating stem cell gene expression. Calcium pathway components are coloured green and planar cell polarity proteins are blue. **(D)** RT-qPCR was performed to compare the expression of the indicated stemness genes in RNA extracted from spheroids incubated for 7 days in suspension culture compared to cells in adherent culture. Gene knock outs are indicated for each and coloured green for Ca pathway and blue for planar cell polarity. Data from three to six biological samples are shown. Means were compared using one-way ANOVA (*p<0.05, **p<0.01, ***p<0.001, ****p<0.0001, error bars are ±1 SEM).

### WNT8B and WNT9B signaling contributes to cancer cell survival and spread in mouse xenograft models

To investigate how WNT8B and WNT9B signaling influences cancer progression *in vivo*, we xenografted female NOD-SCID mice with either OVCAR8 sgRNA control cells or *WNT* DKO cells via intraperitoneal injection. To model dormant residual disease, we euthanized and necropsied mice two weeks post-injection (**Figure 6A**). No macroscopic tumor lesions were detected at this early time point following engraftment (**Figure 6B**), consistent with our previous observations of disease state at this time point (Perampalam et al, 2024a). However, rinsing the abdominal cavities of mice with saline recovered disseminated clusters of cancer cells that resemble residual dormant spheroids, which persist after debulking surgery and chemotherapy in patients. **Figure 6C** shows representative images of reattached spheroids isolated from the abdominal wash of a single mouse from either the OVCAR8 control or the *WNT* DKO xenografted group, with these spheroids being subsequently fixed and stained with Crystal Violet dye. Fewer and smaller spheroids were recovered from *WNT* DKO engrafted mice compared to controls. Considering that mouse cells might also contribute to these spheroids, we additionally quantified human cancer cells in abdominal washes using qPCR to detect human Alu repeats (**Figure 6D**). Our findings demonstrated a substantial decrease in cancer cell abundance in *WNT* DKO cell xenografts after two weeks, indicating that WNT8B- and WNT9B-mediated signaling contributes to the survival of cells at this dormant stage of dissemination.

**Figure 6.**
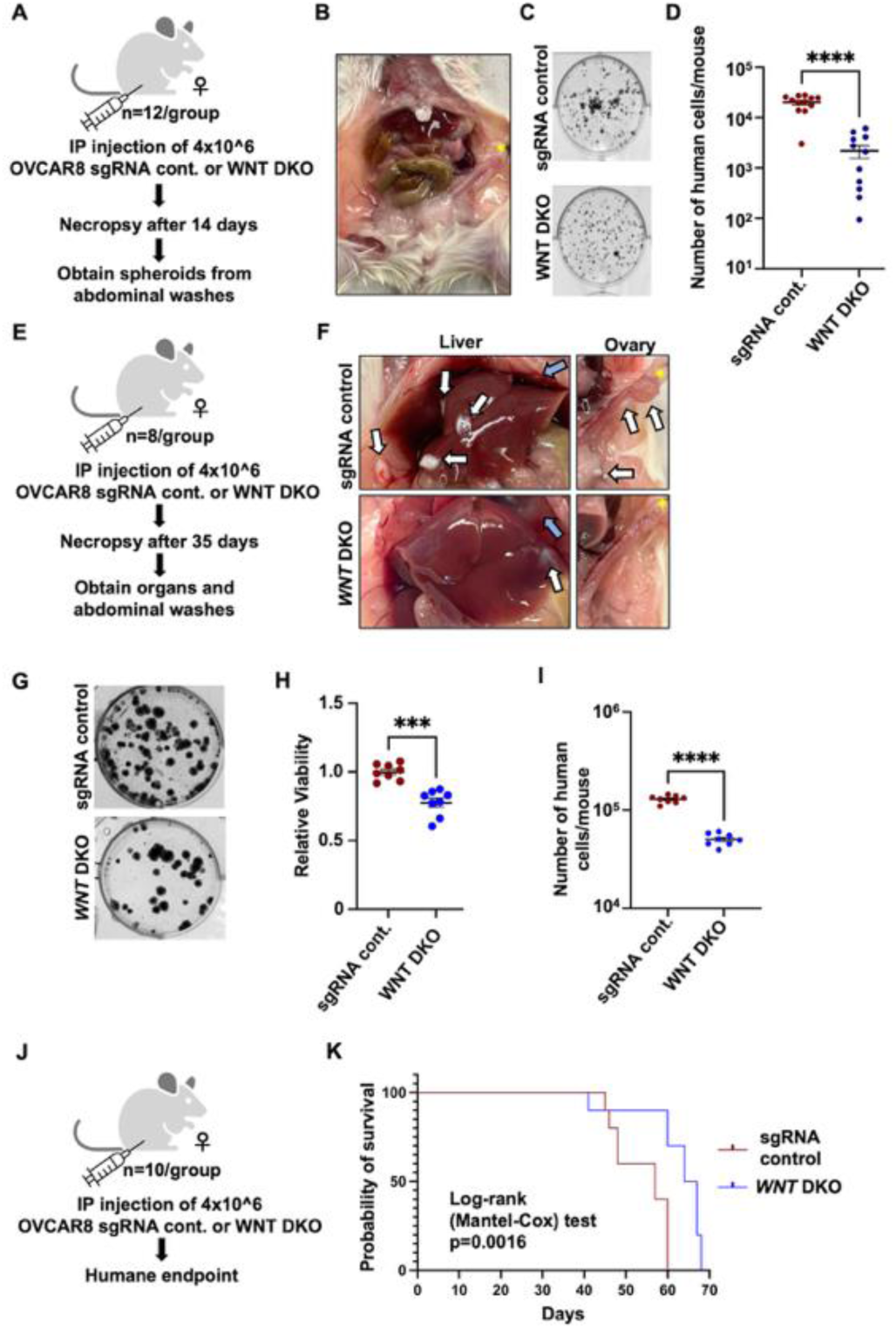
WNT8B and WNT9B deficiency inhibits peritoneal metastasis in xenografted mice. **(A)** Mice were injected intraperitoneally (IP) with either OVCAR8 sgRNA non-targeting control or WNT DKO cells as specified. Mice were monitored daily and euthanized at 14 days post-engraftment for necroscopic analyses. The abdominal cavities of mice were washed with 1 ml PBS to recover disseminated cells. **(B)** Representative image of a xenografted mouse captured during necropsy. An ovary is denoted by a yellow asterisk *. **(C)** Spheroids isolated from abdominal washes were reattached in adherent culture, fixed and stained with Crystal Violet. Representative images of stained spheroids are shown. **(D)** DNA was isolated from cells obtained from abdominal washes, and qPCR was used to detect human Alu repeats, enabling quantification of the number of human cancer cells. Means were compared by t-test (n=12, ****p<0.0001, error bars are ±1 SEM). **(E)** Mice were injected IP with either OVCAR8 sgRNA non-targeting control or *WNT* DKO cells and monitored daily for 35 days. **(F)** Necropsy analyses and rinsing the abdomens to recover disseminated cancer cells were performed as above. Photographs of necropsy findings in the liver and ovary. Locations of ovaries (marked by *), diaphragm (blue arrow), and tumour nodules (white arrows) are indicated in each case. **(G)** Spheroids isolated from the abdominal washes were reattached to adherent culture plates and stained with Crystal Violet, with representative images shown. **(H)** Relative spheroid cell viability was compared between the xenografted groups by extracting the Crystal Violet dye and reading the absorbance values (n=8, ***p<0.001, determined by t-test, error bars are ±1 SEM). **(I)** Detection of human Alu repeats was performed by qPCR (n=8, ****p<0.0001, determined by t-test, error bars are ±1 SEM). **(J)** Mice were injected IP with either OVCAR8 sgRNA non-targeting control or *WNT* DKO cells and monitored until humane endpoint. **(K)** Kaplan-Meier survival analysis of mice engrafted with the specified cell genotypes. Survival was compared by log-rank test (n=10).

We also xenografted mice and euthanized and necropsied them 35 days post-engraftment (**Figure 6E**). This demonstrated that WNT8B and WNT9B loss results in a reduced dissemination of cancer, as evidenced by fewer solid tumors growing on livers and ovaries (**Figure 6F**). Similarly, we recovered significantly fewer spheroids and human cancer cells from the abdominal washes of mice xenografted with *WNT* DKO cells at 35 days (**Figure 6G, H, I**). Finally, we xenografted mice with control and *WNT* DKO cells and monitored them until they reached humane end points (**Figure 6J**). We observed that engraftment with *WNT* DKO cells resulted in significantly longer survival compared to the OVCAR8 control group that received the same number of cells (**Figure 6K**). Taken together, our findings demonstrate that the deletion of WNT8B and WNT9B compromises the viability of dormant spheroid cells and impedes metastatic spread in a mouse model of HGSC dormancy and dissemination.

### WNT8B and WNT9B deletion increases sensitivity to Carboplatin treatment

Ascites-borne cells are known to be highly resistant to conventional chemotherapy such as Carboplatin that is used to treat HGSC patients (Carmi et al, 2024; Ford et al, 2020). We investigated the role of WNT8B and WNT9B signaling in this context by culturing sgRNA control and *WNT* DKO cells in suspension in the presence of a high range of Carboplatin (**Figure 7A**). *WNT* DKO cells showed greater sensitivity to chemotherapy than OVCAR8 control cells, as evidenced by significantly fewer viable spheroids reattaching after Carboplatin treatment (**Figure 7A**), and lower viability as determined by crystal violet staining (**Figure 7B**).

**Figure 7.**
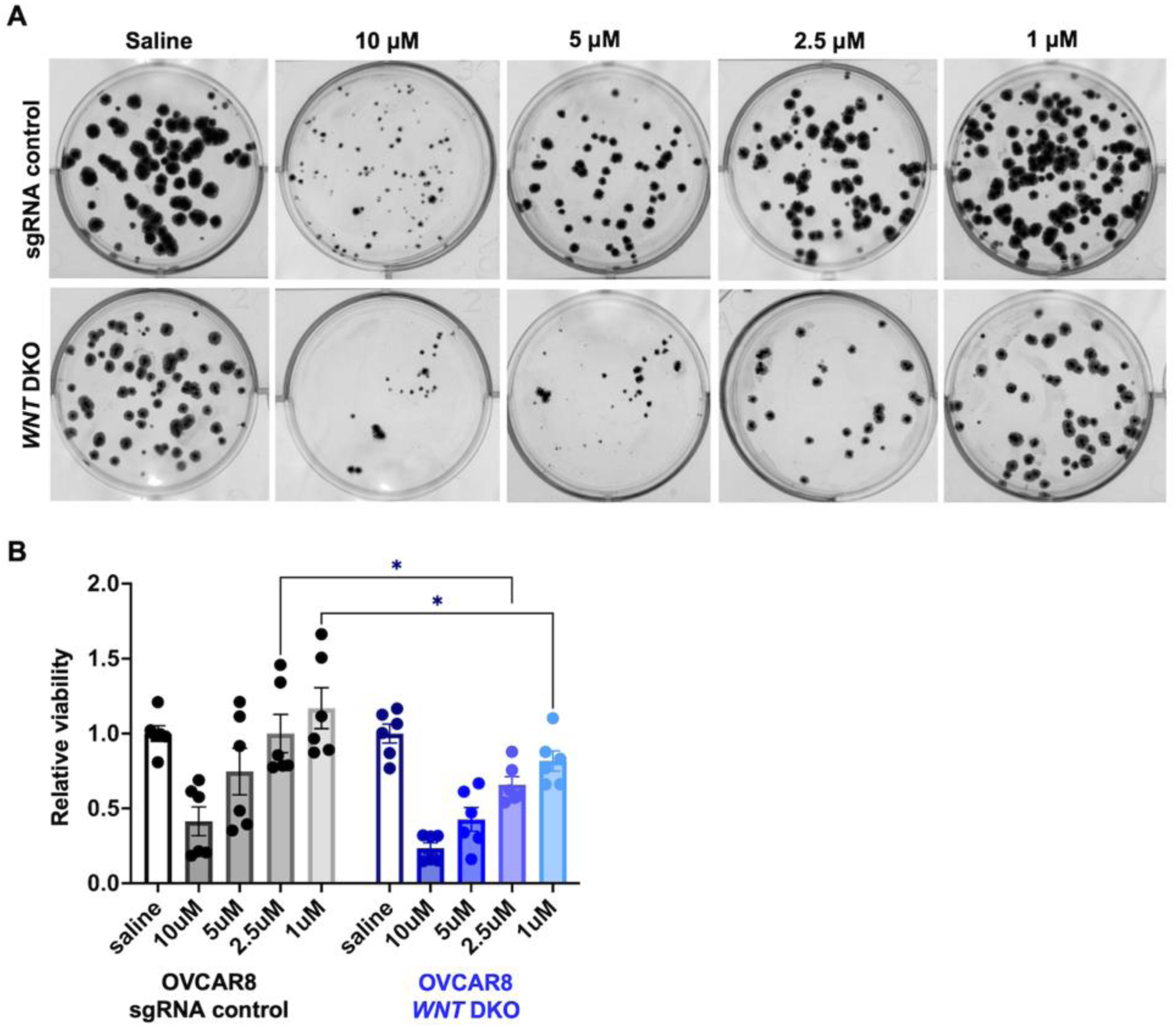
Deletion of WNT8B and WNT9B enhances the effectiveness of chemotherapy. **(A)** OVCAR8 sgRNA non-targeting control, or *WNT* DKO derivatives were seeded in suspension at 600,000 cells per well and cultured with the indicated concentrations of Carboplatin for 7 days. The experiment was conducted in six biological replicates, and representative images of reattached spheroids are shown. **(B)** Re-attached cells were stained with Crystal Violet and retained dye quantity was determined by spectrophotometry. The relative viability of spheroids in each cell genotype was normalised to the survival of the same cell line treated with saline, and statistical comparisons were made using one-way ANOVA (*p<0.05).

Since *in vitro* spheroid culture assays show *WNT8B* and *WNT9B* deletions increase sensitivity of cells to Carboplatin treatment, we next assessed how inhibiting WNT8B and WNT9B signaling affects chemotherapy response in our xenograft model. Mice were engrafted with either OVCAR8 sgRNA control or *WNT* DKO cells (**Figure 8A**). Subsequently, mice were also treated with a low dose of Carboplatin to challenge cells specifically when they are in a dormant state. At 35 days post-engraftment, the mice were euthanized and necropsied. Overall, the OVCAR8 control group exhibited a higher tumor burden, with many more solid nodules attached to the liver, omentum, and ovary, compared to the *WNT* DKO xenografted mice (**Figure 8B**). In addition, fewer viable spheroid cells were isolated from the abdominal washes of mice in the *WNT* DKO treatment group compared to the control cells (**Figure 8C, D**). We also compared the number of human cancer cells recovered using Alu qPCR and it demonstrated that the typical recovery of *WNT* DKO cells at the end of a 35-day experiment was further diminished by Carboplatin (**Figure 8E**). This confirms that the increased sensitivity of *WNT* DKO cells to Carboplatin is also evident *in vivo*.

**Figure 8.**
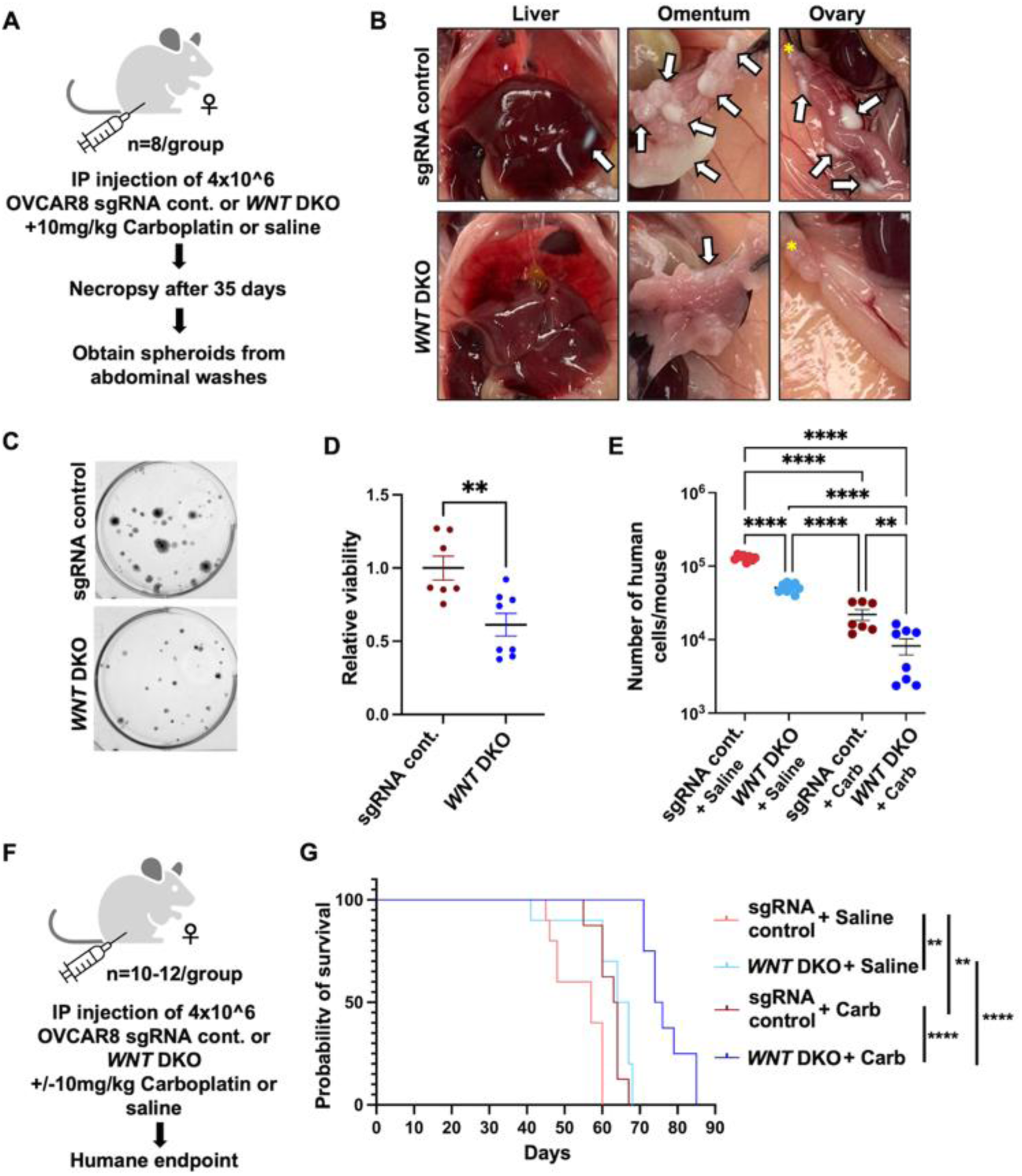
WNT8B and WNT9B loss improves response to Carboplatin treatment in mice. **(A)** Mice were injected intraperitoneally with either OVCAR8 sgRNA non-targeting control or *WNT* DKO cells. Carboplatin was administered by IP injection at a dose of 10 mg/kg, beginning on the same day as cell engraftment, and a second dose one week later. **(B)** At 35 days post-xenograft, mice were euthanized and necropsied. Photographs of necropsy findings in the liver, omentum and ovary were taken. Locations of ovaries (*) and tumour nodules (white arrows) are shown in each case. **(C)** Spheroids collected from the abdominal washes of mice were reattached to adherent culture plates and stained with Crystal Violet. **(D)** The relative viability of spheroids recovered in each group was measured (n=7 mice in OVCAR8 cont. group, n=8 mice in WNT DKO group, **p<0.01, determined by t-test, error bars are ±1 SEM). **(E)** qPCR was conducted to detect human Alu repeats and quantify the number of human cancer cells in the abdominal washes from the indicated genotypes of cells and treatment conditions (**p<0.01, determined by t-test, error bars are ±1 SEM). **(F)** Mice were injected IP with either OVCAR8 control or *WNT* DKO cells. Carboplatin was administered by IP injection at a dose of 10 mg/kg, once per week for the first two weeks. Mice were monitored until humane endpoint. **(G)** Kaplan-Meier survival analysis of mice engrafted with the specified cell genotypes and treated with 10 mg/kg of Carboplatin for two weeks. Survival was compared by log-rank test (****p<0.0001).

To determine the long-term consequences of early Carboplatin treatment on *WNT8B* and *WNT9B* deficient cells when they are dormant, we administered the same dose of Carboplatin to OVCAR8 sgRNA control and *WNT* DKO xenografted mice and monitored disease progression until humane endpoints were reached (**Figure 8F**). Our results demonstrate that blocking WNT8B and WNT9B signaling enhances chemotherapy effectiveness, as evidenced by significantly longer survival in *WNT* DKO engrafted mice treated with Carboplatin compared with both untreated *WNT* DKO and control OVCAR8 and Carboplatin treated cohorts (**Figure 8G**). Overall, our work indicates that a previously unappreciated role for WNT8B and WNT9B in HGSC is relevant for dormant cancer dissemination and resistance to chemotherapy.

## Discussion

In this study, we show that Wnt ligands, WNT8B and WNT9B, identified as top hits in our genome wide dormancy CRISPR screens, play an essential role in HGSC spheroid cells. Previous research suggests that both WNT8B and WNT9B ligands mainly function within the canonical Wnt signaling pathway (Karner et al, 2011; Ngernsombat et al, 2021). However, our results indicate that in the context of HGSC dormant cells, they activate non-canonical Wnt signaling. We found that WNT8B and WNT9B support the survival of spheroids in suspension culture by regulating the expression of *ALDH1A1*, *CD44*, *SOX9*, and other cancer stem cell genes through non-canonical pathways. Inhibition of Wnt signaling through the loss of these ligands results in decreased spheroid cell viability during extended incubation in dormant suspension culture and increased sensitivity of cells to Carboplatin treatment both *in vitro* and *in vivo*. Additionally, *WNT* DKO cells demonstrate a reduced metastatic potential in xenografted mice, as indicated by significantly fewer cancer cells recovered from the abdominal washes and a notable decrease in solid tumor lesions. Our results suggest a novel role for WNT8B and WNT9B in the progression of HGSC, where cancer stem cell properties enhance long term regeneration and survival of cells in spheroids. This reveals that modestly expressed WNT8B and WNT9B are essential mediators of dormancy and dissemination of HGSC and are potential new targets for treating advanced ovarian cancer.

The tumor microenvironment can also influence Wnt ligand-mediated signaling, where these lipid-modified glycoproteins can facilitate long-range signaling (Fang et al, 2024; Kotrbová et al, 2020; Teeuwssen & Fodde, 2019). Exosomal transport could potentially support cancer stem cell maintenance in disseminated spheroids, contributing to metastatic spread. However, our discovery of *WNT8B* and *WNT9B* in multiple pooled knockout CRISPR screens implies a cell-autonomous deficiency effect on viability. If WNT8B and WNT9B ligands mainly signal via paracrine mechanisms in HGSC cells, the heterogeneous composition of spheroids in our screens containing both knockout (for *WNT8B* or *-9B*) and Wnt proficient cells, would be expected to compensate for their loss in rare cells in a single spheroid. Consequently, we would not expect to see such pronounced effects on spheroid viability from the loss of their genes in our pooled CRISPR screens. This indicates that although the tumour microenvironment undoubtedly influences the complexity of Wnt signaling *in vivo*, the intrinsic, cell-autonomous roles of specific Wnt ligands such as WNT8B and WNT9B are significant.

Multicellular spheroids serve as niches for cancer stem cells, which contribute to the acquisition of resistance to conventional therapies (Alizadeh et al, 2025). Canonical Wnt/β-catenin signaling has been widely associated with the maintenance of CSCs in various cancers (Song et al, 2024). Here we show that non-canonical Wnt signaling, driven by WNT8B and WNT9B ligands, activates cancer stem-like programs within HGSC spheroids, thereby aiding their survival during dormancy. Given the previously unrecognised role of these rare Wnt ligands in HGSC cells, the combination of standard chemotherapy, such as Carboplatin, with novel inhibitors of non-canonical Wnt pathways or rare Wnt ligands may offer a promising therapeutic strategy to target cancer stem-like cells and overcome chemoresistance.

The family of Wnt ligands contains 19 members and there is ample evidence that many are expressed in HGSC (Fang et al, 2024; Kotrbová et al, 2020; Piki et al, 2023; Reinartz et al, 2016). Consequently, the loss of function of single or small subsets of ligands may be compensated by others. In our experiments, single or double knock out of WNT8B and −9B was often quite deleterious to viability or stem cell gene expression in HGSC. It is possible that other Wnts may also function in HGSC such as WNT5A, −7A, −10A, and 11 as reported by others and contribute to these functions. However, the power of our screen approach demonstrates that even in as large a family as Wnt ligands, specific ones possess key roles even when not the most highly induced or expressed. This further suggests that Wnt inhibition as a therapeutic approach may not need to broadly inhibit many family members such as through PORCN inhibitors that are associated with toxicity to normal tissues.

In conclusion, our study highlights the important role of non-canonical Wnt signaling in maintaining the survival of spheroid cells in dormancy by regulating cancer stemness programs. We demonstrate that inhibition of Wnt signaling through the loss of WNT8B and WNT9B ligands significantly impedes disease progression in mouse models of metastasis and improves the sensitivity of HGSC cells to chemotherapeutic treatment.

